# Soil-free pathogenicity assay of *Fusarium oxysporum* f.sp. *vasinfectum* race 4 on Pima cotton (*Gossypium barbadense*) seedlings

**DOI:** 10.1101/2021.04.01.438137

**Authors:** Lia D. Murty, Won Bo Shim

## Abstract

One of the devastating early season diseases of cotton is Fusarium wilt caused by *Fusarium oxysporum* f. sp. *vasinfectum* (Fov). Recent emergence of highly virulent Fov race 4 (Fov4) and its aggressiveness toward *Gossypium barbadense* (pima) cultivars are raising significant concerns for the US cotton industry. One of the key challenges in studying Fov4 virulence and cotton Fusarium wilt pathogenesis is establishing a disease assay strategy that can help researcher overcome several technical challenges, including efficient infection and highly reproducible and consistent symptom development. Here, we have developed a small-scale, soil-free Fusarium wilt disease assay that can complement conventional assays with faster symptom development and high reproducibility in infected pima cotton seedlings. Our data showed statistically significant differences (p<0.0001) between Fov4-infected and non-infected pima cotton at 4 and 6 days post inoculation (dpi) when compared to control experiments. At 6 dpi, longitudinal observations under magnification showed Fov4 colonization in primary xylem of infected plants, which is a common symptom observed in Fov4 triggered Fusarium wilt in pima cotton. While this is an artificial assay system, this soil-free disease testing strategy can offer another strategy to supplement current assays when studying pathogen-host interaction in soil-borne diseases.

## INTRODUCTION

Cotton is an important globally traded commodity worldwide with primary use in the textile and clothing industry. Cotton is recognized as a major cash crop around the world, and the socio-economic importance particularly in developing economies is well recognized. In the US, cotton (*Gossypium* spp) is cultivated in many southern states, with Texas as the top cotton producing state with more than 40% of the total US production. Majority of cotton species cultivated in the US are upland cotton varieties (*G. hirsutum*), with about 3% of the US production being Pima cotton (*G. barbadense*), a finer and higher value fiber, mostly grown in California, Arizona, and west Texas. One of the key early season diseases of cotton is Fusarium wilt caused by *Fusarium oxysporum* f. sp. *vasinfectum* (Fov) (Skovgaard et al. 2001). In the US, race 1 (Fov1) is known to be the predominant pathogen especially in upland cotton producing fields. It is important to note that Fov1 requires root-knot nematodes (RKN) to infect cotton and thus minimizing disease outbreaks with nematicides or RKN resistant cotton varieties has been largely successful.

Significantly, a highly virulent race 4 (Fov4), originating in India, was identified in the California San Joaquin Valley in 2004 (Kim et al. 2005), west Texas in 2017 (Bell et al. 2019) and New Mexico in 2020 (Zhu et al. 2020). Fov4 was determined as the pathogen responsible for dead seedlings and black streaks inside tap roots of wilting Pima cotton plants. The detection of Fov4 in the southwestern US has justifiably caused alarm in the US cotton industry. Similar to Fov1, Fov4 colonizes roots and vascular system resulting in discoloration, wilting and death. Due to its seed-borne and soil-borne characteristics, Fov4 can be transmitted via seeds and on equipment, raising concerns over containment (Liu et al. 2011). Fov4 is now considered an endemic pathogen in California while the presence in Texas is recent and remains spatially isolated. Geographically, Texas is also unique as this location is where western Pima production transitions to upland cotton production to the east. Though upland cotton was once thought to be less susceptible to Fov4, we are now learning that Fov4 may be an inoculum density-dependent disease and can pose serious threat to upland cotton production as well (Bell et al. 2019).

Since the emergence of Fov4 in the US, a number of studies were performed to improve pathogen detection, population diversity, and distribution surveys (Bell et al. 2019; Cianchetta et al. 2015; Doan 2014; Ortiz et al. 2017). In addition, several disease assay methods were described to test Fov4 virulence against cotton varieties (Cianchetta et al. 2015; Ortiz et al. 2017; Wang et al. 2018). While there are some modifications, majority of these methods are based on conventional root-cup dip method (Smith et al. 1994) or pathogen spore injection into pots where cotton seedling were grown (Ortiz et al. 2017). Disease severity was typically assessed by rating symptoms visually observed on above surface cotton leaves, e.g. growth suppression, yellowing and wilting. While we tried to apply these disease assay strategies to our Fov4-cotton interaction research, we faced two major challenges. One was inconsistencies in infection and disease development when inoculating Fov4 spore suspension into cotton seedling pots (Ortiz et al. 2017). The other limitation was that we could not investigate infection process occurring at the Fov4-cotton seedling interface to further understand Fov4 virulence at the molecular and cellular level.

To overcome these limitations, we developed a soil-free Fusarium wilt assay where we can infect Pima cotton with high efficiency while visually observing cotton seedling wilt and rot in a petri dish. The method described here can easily be applied to multiple soilborne pathogens and its plant hosts. Results from this study show this soil-free pathogenicity assay using the Fov4-cotton was highly reproducible and efficient at producing symptoms while requiring minimal equipment. Fov4-cotton samples can be further used in microscopic observation as well as in nucleic acid or protein extraction for molecular and cellular host-pathogen interactions research.

## MATERIALS AND METHODS

### Fov4 inoculum preparation

*Fusarium oxysporum* f. sp. *vasinfectum* race 4 (Fov4) strain used in this study was isolated from diseased Pima cotton plants acquired from El, Paso, Texas (Courtesy of Dr. Tom Isakeit, Texas A&M AgriLife Extension). Identification of this Fov4 isolate was confirmed using the method described in Doan, et. al. 2014 and AmplifyRP^®^ Acceler8^®^ for Fov4 rapid DNA kit (Product No. ACS 19700/0008) (Doan 2014). Fov4 inoculum was prepared by growing on ISP2 agar medium (HIMEDIA 2019) and flooding 7-14 day cultures with sterile water and plastic spreader to suspend conidia into solution, then solution was filtered through double layered sterile miracloth (Bell et al. 2019). Conidia concentration was determined by hemocytometer.

Steel cut oats (SCO) medium was prepared by adding 20 g of steel cut oats and 20 ml deionized water into a 100-ml flask. These flasks were gently shaken several times and autoclaved for 30 min. When cooled to room temperature, these flasks were vigorously shaken to breakup clustered SCO. Flasks were stored at room temperature for 24 h, and autoclaved again for 30 min. Fov4 SCO inoculum was prepared by inoculating a 1-ml conidia suspension (1×10^4^ conidia/ml) to SCO medium followed by incubation at 28°C for 5 days, shaken daily to promote fungal dispersal and uniform inoculum growth. Prepared Fov4 SCO inoculum can be stored at 4°C up to 14 days until use. As controls, we used maize pathogen *F. verticillioides* and saprophytic *Penicillium* species on pima cotton seedlings following the procedure described above.

### Pima seedling preparation

Pima cotton (PhytoGen^®^, No. PHY841RF; courtesy of Dr. Thomas Chappell, Texas A&M University) were surface sterilized with 10% bleach and 70% ethanol following standard laboratory practice. Seeds were placed on a sterile cotton round in a group of three and covered with a second cotton round which was then applied with 10 ml of sterile water. Two cotton round sandwiches were placed in a plastic sandwich bag, subsequently enclosed and taped to minimize moisture loss but not completely sealed to permit airflow. These plastic bags were then taped to a laboratory window with ample day light or in a growth chamber, and allowed to germinate and grow for up to 15 days. For our soil-free pathogenicity assay, we selected cotton seedlings that developed cotyledon leaves with primary root-shoot length between 5-7cm. We eliminated any seedling that exhibited visible blackening at the root shoot junction (RSJ; anticipated as the soil line) and into the roots likely caused by seed-borne pathogen contamination. As a negative control, we inoculated Fov4 on soybean seedlings following the procedure described above.

### Pathogenicity Assay

Selected cotton seedlings were placed on to new sterile half cotton rounds (1/2 circle) (Fig 1A). Two seedlings were placed in a standard petri dish, and the roots covered with approximately 1 g of SCO inoculum (Fig 1B). Autoclaved SCO was used as control (CT). Subsequently, a quarter (1/4) cotton round was placed on top of cotton seedling and SCO (Fig 1C). Lastly, the petri dish lid was placed on top with two small pieces of tap to hold top in place. Assembled plates were placed under 12 hr light/dark cycle at room temperature (22∼24°C). Fusarium wilt symptoms were monitored daily, and two replicates (1 plate) for each group was terminated at 4 dpi to observe any signs of Fusarium wilt symptom development. The remaining replicates were terminated at 6dpi for final assessment and vertical cuts of RSJ observation under 4×0.1 magnification. Ten replicates were used for each treatment: Fov4 infected on pima seedlings and autoclaved SCO on pima seedlings.

**Figure 1.**
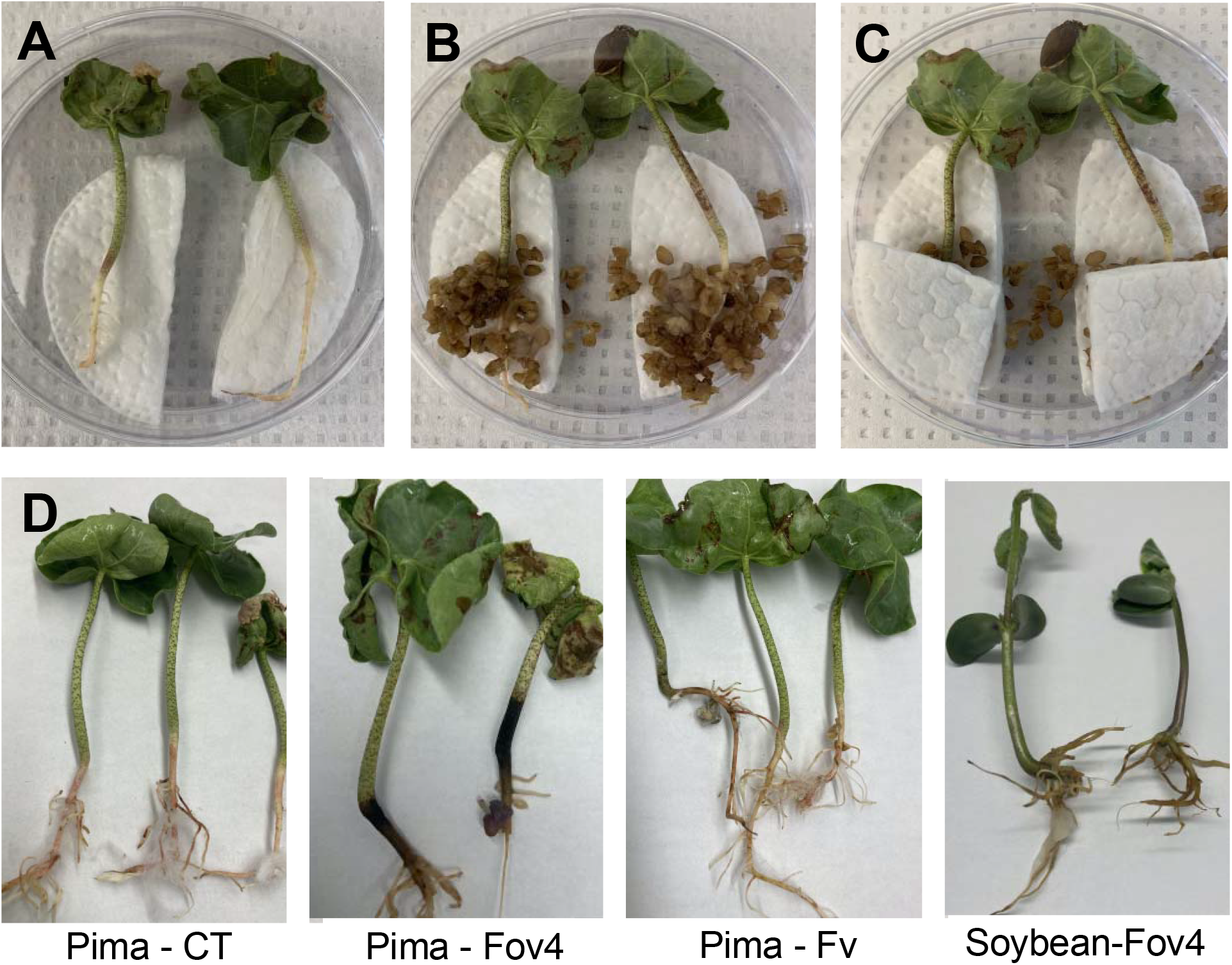
Soil-free Fusarium wilt assay setup in a standard petri dish. (A) Cotton seedlings identified for use in pathogenicity assay displaying no symptoms of infection, (B) Cotton seedlings with approximately 1g of steel cut oat inoculum, (C) Final covering of cotton roots with steel cut oat inoculum. (D) Fusarium wilt assay perform with pima – negative control (autoclaved oatmeal: CT), pima – Fov4, pima – *F. verticillioides* (Fv), and soybean – Fov4.

### Disease Rating and Assessment

We followed disease symptoms for 5 days, and on day 6 cotton seedlings were removed from petri dish for examination under a microscope. Ratings were on a scale from 0 to 5, and the visual representation of our disease ratings is shown in Figure 2 and Table 1. Disease ratings were based on the symptoms observed at and above the RSJ. The coloring at the RSJ and along the shoot was used to assess disease. Additionally, the visibility of gossypol glands was used as well. Disease ratings were statistically analyzed using Graphpad Prism (San Diego, CA) using a one-way ANOVA followed by a Tukey post hoc test with 95% confidence interval.

**Figure 2.**
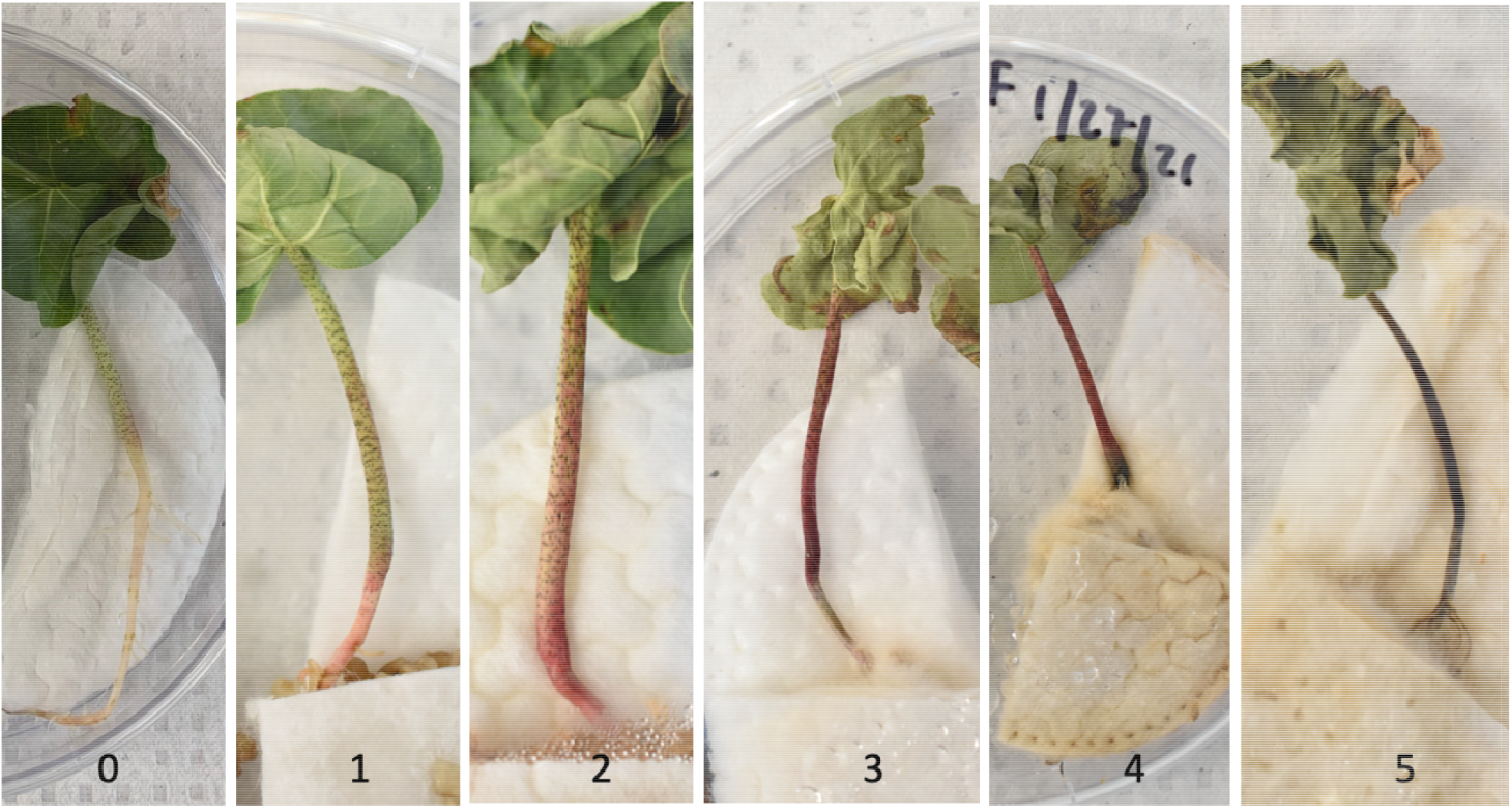
Disease progression with rating system used to assess pathogenicity assay. The disease rating follows the progression of pink pigment starting at the base of the shoot to a change to black discoloration through the entire shoot.

**Table 1.** Criteria used in the disease rating assessment of Fov4 infected pima cotton seedlings.

| Rating | Root Shoot Junction (RSJ) Color | Shoot Color | Visible gossypol glands on shoot |
| --- | --- | --- | --- |
| 0 | Green | Green | visible entire length |
| 1 | Light Pink | Green | visible entire length |
| 2 | Dark pink | Light pink | visible entire length |
| 3 | Dark purple | Dark purple | visible middle of shoot up to leaves |
| 4 | Black | Dark purple | visible only top of shoot under leaves |
| 5 | Black | Black | not visible |

## RESULTS and DISCUSSION

Two treatments, Fov4-inoculated (Fov4) seedlings and negative control (CT) seedlings, were concurrently prepared and monitored daily. We also tested multiple control experiments, *i*.*e*. Fov4-soybean, *F. verticillioides*-pima, *Penicillium*-pima, and confirmed that the wilt symptom we observe in Fov4-pima is specific to this pathogen and its host plant (Fig 1D). Treatments showed no statistical significance at 2dpi. On 4dpi and 6dpi negative controls compared to Fov4 infected treatment showed p<0.0001. We started observing signs of stress in both Fov4 and CT seedlings two days post inoculation (dpi). However, we could clearly distinguish two treatments since CT seedlings showed minimal pink pigmentation. Notably, Fov4-treated seedlings gradually exhibited pink pigment progressing up from the base of the roots to the entire stem by 6 dpi. At 6 dpi, the RSJ rotting was clearly visible with black discoloration. Two replicates from each treatment were terminated at 4 dpi, and longitudinal cross sections of RSJ were examined under a microscope at 4×0.1 magnification to observe symptom development. We did not see distinguishable differences in cross sections. However, pigment accumulation on seedling surface tissues were clearly recognizable with brown, darkening pink to violet pigmentation in Fov4 plants. At 6 dpi all replicates were excised from Fov4 SCO inoculum and CT SCO inoculum. We followed our disease rating system (Fig 2, Table 1) to assess the severity of Fusarium wilt and results are shown in Figure 4. In Fov4-infected samples, excised plants maintained some structural root stability while still showing symptoms of Fov4 infection. When plants were excised, we were able to see that the rotting had not completely disintegrated the roots. Longitudinal section of the root shoot junction under microscope examination showed that at 6 dpi, the difference between CT and Fov4 seedlings were clearly noticeable upon examination of the vascular tissue. Figure 3 shows the difference in root vascular tissue; the root of Fov4 seedlings clearly showing necrotic symptom in the vascular tissues in contrast to CT seedlings.

**Figure 3.**
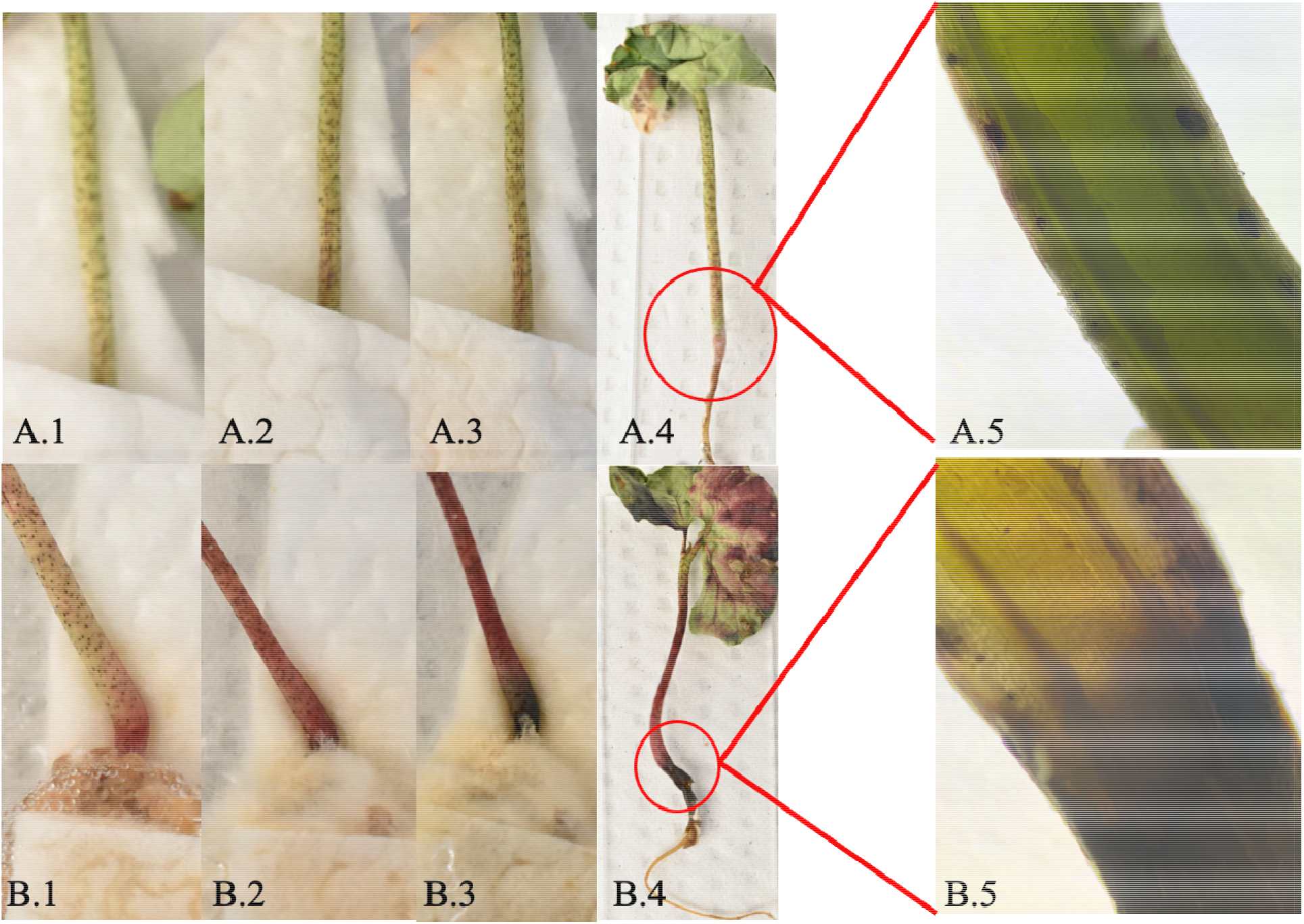
Fov4 infection of pima cotton over six days post inoculation. **A**.**1** Control day 2 dpi, **A**.**2**. Control day 4 dpi, **A**.**3**. Control 6 dpi, **A**.**4**. Excised control replicate at day 6, **A**.**5**. Magnification (4×0.10) of root shoot junction (RSJ) of control plant 6 dpi; **B**.**1** Fov4 infection 2 days post inoculation, **B**.**2**. Fov4 infection 4 dpi, **B**.**3**. Fov4 infection after 6 dpi, **B**.**4**. Excised infected plant 6 dpi, **B**.**5**. Magnification (4×0.10) of RSJ of 6 dpi Fov4-infected cotton.

**Figure 4.**
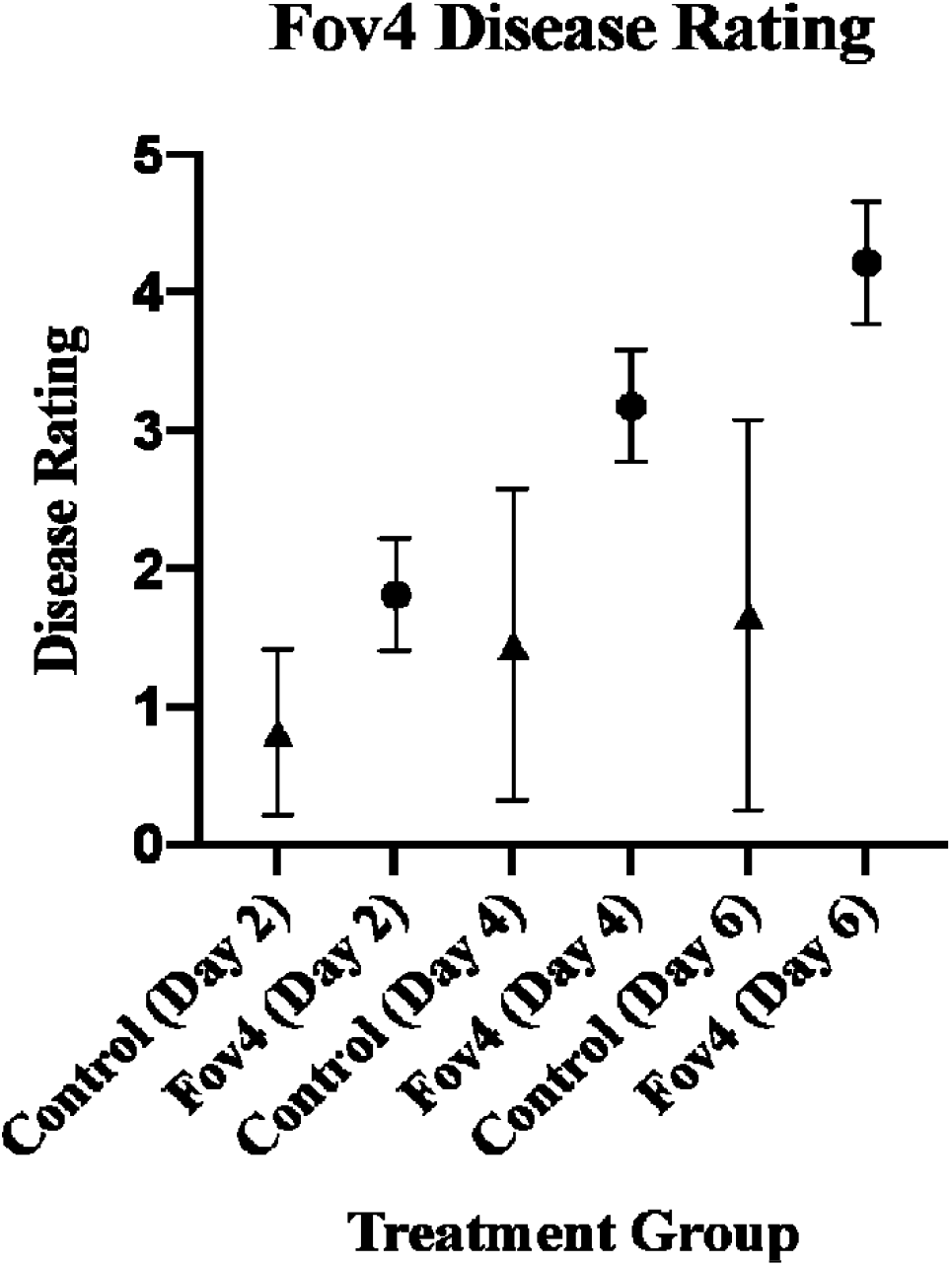
Fusarium wilt disease rating progression compared in Fov4-infected (Fov) and negative control (Control) samples over six days. Disease ratings were statistically analyzed using a one-way ANOVA followed by a Tukey post hoc test with 95% confidence interval.

In this study, our aim was to design and test a pathogenicity assay system capable of overcoming the most challenging aspects of studying Fusarium wilt of cotton caused by Fov4. As described Bell, et. al. 2017, Fov4 does not require root-knot nematode for cotton infection unlike Fov1 (Bell et al. 2017). However, it is also interesting to note that direct wounding and injection of Fov4 inoculum into cotton seedlings does not also provide reliable symptom development (Liu et al. 2011). Therefore, current Fov4 pathogenicity assay relies on indirect fungal inoculum injection into soil where cotton seedlings are grown, and this practice can lead to inconsistent and sometime non-reproducible assay results (Bell et al. 2019; Bell et al. 2017). With our soil-free assay system, we were able to achieve high reproducibility among treatment groups. In other soil pathogenicity assays, the assessment of disease was mostly done on above ground parts by measuring shoot growth suppression, leaf yellowing and overall plant wilting (Bell et al. 2019; Bell et al. 2017; Liu et al. 2011). Here we were able to monitor above-ground disease symptom development but also examine root system when experiment was terminated. If needed, seedling samples can be removed from petri dish for root infection examination and place back into assay setting to continue the infection progress. However, this may require stringent sanitation practice and caution to minimize contamination and plant seedling wounding that can lead to inaccurate interpretation.

Another key advantage of this assay system is the efficiency and minimal material requirement. We were able to achieve consistent and reliable disease assay in 5 days, without the use of soil, pots, fertilizer, and growth chambers. Symptom development of Fov4 infection in conventional soil pathogenicity assays has been evaluated 38 days post inoculation (Bell et al. 2019). As described earlier, some of the recent modified Fusarium wilt assays required large amount of pathogen inoculum, transplanting of cotton seedlings, and inefficient inoculation strategies (Cianchetta et al. 2015; Ortiz et al. 2017; Smith et al. 1994; Wang et al. 2018). The use of SCO provided a uniform Fov4 inoculation with minimal material requirements. There was also no need for weekly fertilization of plants since we were able to observe Fusarium wilt symptoms in less than 7 dpi. By starting the pathogenicity assay when shoot lengths were 5-7cm allowed us to eliminate any plant material contaminated with various seed-borne pathogens, including Fov4. Reproducibility was evident by low coefficient of variation among the disease ratings in each of the Fov4 treatment groups, at 2 dpi there was only 22% variation among replicates, 13% at 4 dpi, and 10% at 6 dpi. While re-isolation of pathogen has been done in other pathogenicity assays, the experimental set up was able to be conducted in biosafety cabinets under sterile conditions and until termination.

In summary, we developed a soil-free Fusarium wilt assay where we can infect Pima cotton with Fov4 with high efficiency and consistency while visually observing cotton seedling with wilt and rot symptoms in a petri dish. We acknowledge that this is an artificial assay system that does not truly reflect field conditions. But in some regard, similar criticism can be made about conventional plant disease assays where autoclaved soil in pots are maintained in growth chambers or green house settings. Here, we make an argument that this soil-free assay system can provide a different perspective when studying pathogen-host interaction in soil-borne diseases. This method can easily be applied to a variety of soilborne pathogens and its plant hosts, especially with soil-borne *Fusarium* species. Furthermore, this soil-free plant disease assay system can be used in K-12 science laboratories to educate students about key plant pathology concepts, e.g. infection mechanism, host specificity, non-host immunity, with minimal equipment and facility needs.

## ACKNOWLEDGEMENTS

The research was supported by Texas A&M T3: Triads for Transformation Program and Texas A&M AgriLife Research Strategic Initiative.

